# Spatially Resolved Estimation of Metabolic Oxygen Consumption From Optical Measurements in Cortex

**DOI:** 10.1101/827865

**Authors:** Marte J. Sætra, Andreas V. Solbrå, Anna Devor, Anders M. Dale, Gaute T. Einevoll

**Affiliations:** Centre for Integrative Neuroplasticity, University of Oslo, Oslo, Norway; Department of Physics, University of Oslo, Oslo, Norway; Department of Radiology, University of California San Diego, La Jolla, CA, United States; Department of Neurosciences, University of California San Diego, La Jolla, CA, United States; Martinos Center for Biomedical Imaging, MGH, Harvard Medical School, Charlestown, MA, United States; Faculty of Science and Technology, Norwegian University of Life Sciences, Ås, Norway

**Keywords:** CMRO_2_, cortex, metabolism, analysis, estimation

## Abstract

The *cerebral metabolic rate of oxygen (CMRO*_2_) is an important indicator of brain function and pathology. Knowledge about its magnitude is also required for proper interpretation of the blood oxygenation level dependent (BOLD) signal measured with functional MRI (fMRI). The ability to measure CMRO_2_ with high spatial and temporal accuracy is thus highly desired. Traditionally the estimation of CMRO_2_ has been pursued with somewhat indirect approaches combining several different types of measurements with mathematical modeling of the underlying physiological processes. Given the numerous assumptions involved, questions have thus been raised about the accuracy of the resulting CMRO_2_ estimates. The recent ability to measure the level of oxygen (pO_2_) in cortex with high spatial resolution in *in vivo* conditions has provided a more direct way for estimating CMRO_2_. CMRO_2_ and pO_2_ are related via the Poisson partial differential equation. Assuming a constant CMRO_2_ and cylindrical symmetry around the blood vessel providing the oxygen, the so-called Krogh-Erlang formula relating the spatial pO_2_ profile to a constant CMRO_2_ value can be derived. This Krogh-Erlang formula has previously been used to estimate the average CMRO_2_ close to cortical blood vessels based on pO_2_ measurements in rats.

Here we introduce a new method, the *Laplace method*, to provide spatial maps of CMRO_2_ based on the same measured pO_2_ profiles. The method has two key steps: First the measured pO_2_ profiles are spatially smoothed to reduce effects of spatial noise in the measurements. Next, the Laplace operator (a double spatial derivative) in two spatial dimensions is applied on the smoothed pO_2_ profiles to obtain spatially resolved CMRO_2_ estimates. The smoothing introduces a *bias*, and a balance must be found where the effects of the noise are sufficiently reduced without introducing too much bias. In this model-based study we explore this balance in situations where the ground truth, that is, spatial profile of CMRO_2_ is preset and thus known, and the corresponding pO_2_ profiles are found by solving the Poisson equation, either numerically or by taking advantage of the Krogh-Erlang formula. MATLAB code for using the Laplace method is provided.

## 1. Introduction

The level of consumption of oxygen by metabolic processes, that is, the *cerebral metabolic rate of oxygen (CMRO*_2_), is an important indicator of brain function and pathology. Further, knowledge about the magnitude of the CMRO_2_ is also required for a proper interpretation of the blood oxygenation level dependent (BOLD) signal measured in functional MRI (fMRI) studies [Buxton, 2010]. The ability to measure CMRO_2_ with high spatial and temporal resolution in cortex is thus crucial. Traditionally the CMRO_2_ has been estimated from several different types of measurements combined with mathematical modeling of the underlying physiological processes [Buxton, 2010]. Given the numerous assumptions and experimental limitations typically involved, questions have been raised about the accuracy of the estimates of the CMRO_2_ provided by these complex and somewhat indirect approaches [Sakadžić et al., 2016].

The possibility to optically measure the partial pressure of oxygen (pO_2_) around cortical blood vessels with high spatial resolution *in vivo* [Sakadžić et al., 2010] has provided a more direct way to estimate the CMRO_2_. In Sakadžić et al. [2016] they used measured pO_2_ profiles around arterioles in rats to estimate the average CMRO_2_ in the vessel’s vicinity, that is, within a radius of ∼100 *µ*m. They based their estimates on the Krogh-Erlang formula relating the pO_2_ to the CMRO_2_ in a cylinder section around an arteriole providing the brain tissue with oxygen [Krogh, 1919; Goldman, 2008].

The fundamental equation relating the pO_2_ and the CMRO_2_ is the Poisson equation

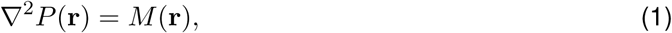

where *P* (**r**) represents pO_2_, and *M* (**r**) is a measure of the local CMRO_2_. The Krogh-Erlang formula (Equation 6) gives the solution to this partial differential equation, that is, the radial profile of *P*, for the particular case where (i) the CMRO_2_ (*M* (**r**)) is assumed to be a constant, and (ii) all the oxygen provided by the center arteriole is assumed to be consumed within a radial basin with radius *R*_*t*_. In Sakadžić et al. [2016], experimentally measured pO_2_ profiles were fitted to this formula to provide estimates for *M* (and thus CMRO_2_).

The approach of Sakadžić et al. [2016] is *global* in the sense that it fits the entire measured profile *P* (**r**) to the Krogh-Erlang formula to obtain an estimate for the assumed constant value of *M*. A more direct way to estimate *M* (**r**) from Equation 1, is to apply the Laplace operator ∇^2^ directly to the measured *P* (**r**) to obtain a *local* measure of *M* (**r**). Unlike the Krogh-Erlang model approach, this *Laplace approach* will provide a spatially resolved map of CMRO_2_ estimates around the arterioles, based on the same pO_2_ measurements. The method is thus not restricted to estimating an assumed constant value of *M*. Further, the Laplace method is not restricted to situations with radially symmetric pO_2_ profiles as when a single arteriole provides all oxygen. The development and testing of the Laplace method are the topics of the present paper.

The double spatial derivatives in the Laplace operator make this Laplace method inherently very sensitive to noise in the measured spatial pO_2_ profiles. In order to have a practical method for CMRO_2_ estimation, the pO_2_ profiles must thus be spatially smoothed to reduce the effects of the noise. Smoothing introduces a *bias*, that is, a systematic error in the estimates, and a balance must be found where the effects of the noise are sufficiently reduced without introducing too much bias. In the present model-based study we explore this balance by examining the accuracy of CMRO_2_ estimates in situations where the ground truth, that is, spatial profile of *M* (**r**) is preset and thus known, and the corresponding profiles *P* (**r**) are found by solving Equation 1, either numerically or by taking advantage of the Krogh-Erlang formula.

The manuscript is organized as follows: In Section 2 we describe the Laplace method, the methods used to provide model-based pO_2_ profiles used in the testing, and the metrics used to quantify the accuracy of the resulting estimates. In Section 3 we first illustrate the method and the necessary compromise between reducing noise and limiting bias when choosing the level of spatial smoothing. Next, we systematically explore the accuracy of CMRO_2_ estimates for a variety of situations with different levels of noise, different grid sizes of the pO_2_ measurement, and different levels of smoothing. In these systematic explorations of the efficacy of the method, the simple single-arteriole situation where the Krogh-Erlang formula gives the ground truth, is considered for simplicity. Later, we illustrate the use of the Laplace method on more complicated situations where several arterioles provide the consumed oxygen, or the CMRO_2_ varies with position. In Section 4 we discuss the Laplace method and its further development and use.

## 2. Methods

### 2.1. Forward modeling of oxygen consumption

The blood-tissue O_2_ transport is assumed to be caused by diffusion and is described mathematically by the Poisson equation. Under steady-state conditions, that is, no time dependence of the oxygen partial pressure *P*, the relationship between this pressure and the net rate of oxygen consumption *s*(**r**) in the tissue can be described by [Goldman, 2008; Sakadžić et al., 2016]:

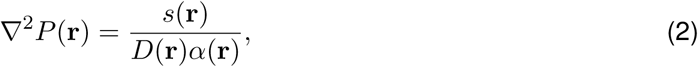

where ∇^2^ is the Laplace operator, *D*(**r**) is the diffusivity, and *α*(**r**) is the solubility of the medium. The equation can be written more compactly as

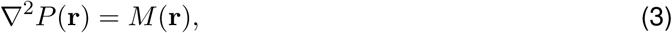

where

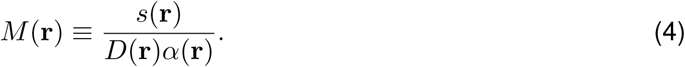

Here, *M* (**r**) is a new position-dependent variable encapsulating the oxygen consumption in the neural tissue.

By introducing a characteristic length *r*^*^ and a characteristic oxygen consumption *M*^*^, we can rewrite Equation 3 in a dimensionless form which is useful in the further analysis:

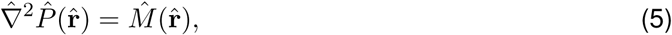

where 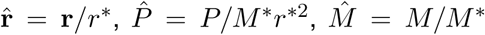, and 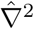 is the Laplace operator in terms of the dimensionless position variables. In this dimensionless form, the number of model parameters is effectively reduced by one, making the further analysis simpler.

Equation 3, and the dimensionless version in Equation 5, in principle describe the spatial profile of the oxygen pressure for any set of oxygen sinks (metabolic consumption, *M* > 0) and sources (oxygen provided by vessels, *M* < 0). The variable *M* (**r**) describes the net oxygen consumption, that is, the difference between oxygen sinks and sources at position **r**.

In general, both the oxygen pressure *P* (**r**) and the net oxygen consumption *M* (**r**) depend on the position in three-dimensional space. However, in the present application we assume no axial diffusion of oxygen, that is, no diffusion in the direction parallel to the blood vessel providing the oxygen. Thus *P* (**r**) = *P* (*x, y*) and *M* (**r**) = *M* (*x, y*).

#### 2.1.1. Krogh-Erlang model

In the well-known Krogh-Erlang model [Krogh, 1919], a cylindrical geometry, mimicking a straight segment of a blood vessel, was used to model the metabolic consumption of oxygen provided by capillaries in muscles. In Sakadžić et al. [2016], the same model was used to study metabolic consumption of oxygen provided by arterioles in brain tissue. The model describes the blood vessel as a small cylinder with radius *R*_ves_ supplying a tissue cylinder with radius *R*_t_ with oxygen. The further assumptions are (i) uniform consumption of oxygen in the tissue, that is, constant *M* outside the vessel, (ii) no axial diffusion of oxygen, (iii) *P* = *P*_ves_ at *R*_ves_, and (iv) no pressure gradient at the surface of the tissue cylinder, that is, *dP/dr* = 0 at *R*_t_. With these four assumptions, the solution of Equation 3 is found to be

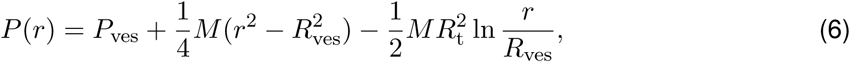

for *R*_t_ ≥ *r* ≥ *R*_ves_. This so-called Krogh-Erlang formula predicts the oxygen pressure *P* in the tissue as a function of the distance *r* from the vessel’s center. For our application we set *P* (*r*) = *P*_ves_ if *r* < *R*_ves_.

Equation 6 can be written in dimensionless form as

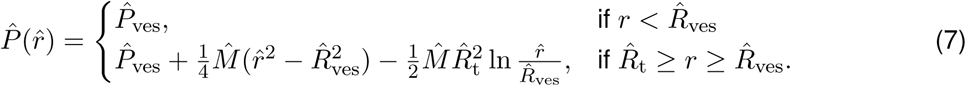

Here we also have introduced 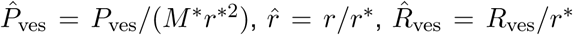 and 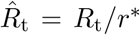 Further, the boundary condition 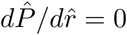 for 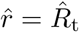 is assumed.

#### 2.1.2. FEniCS model

The Krogh-Erlang formula relates the oxygen consumption and the partial oxygen pressure under very specific conditions. Another option is to solve Equation 5 numerically. This allows for the solutions for more general cases, such as a more complicated geometry with, for example, several arterioles providing oxygen, or a variable oxygen consumption. We implemented Equation 5 in the finite element software package FEniCS [Logg et al., 2012], and verified the implementation by comparing the result to that of the Krogh-Erlang formula.

The FEniCS implementation solves the variational formulation of Equation 5: Let *V* be a space of test functions {*v*_1_, *… v*_*N*_} on the computational domain Ω. We aim to find 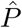 such that

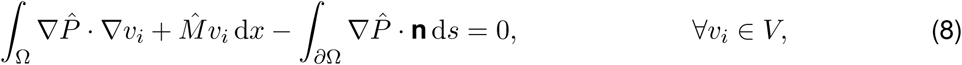

where *∂*Ω denotes the boundary of the domain, and **n** is a normal vector pointing out of the domain. This variational form is obtained by multiplying Equation 5 with the test function *v*_*i*_ and integrating over Ω, followed by integration by parts of the Laplacian term. Note that as we apply a fixed value for 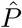 by the blood vessel and no pressure gradient at the boundary of the domain, the boundary integral in Equation 8 vanishes.

#### 2.1.3. Noise

We add additive Gaussian noise to the test data using the `normrnd` function in MATLAB. For each value 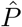 of oxygen partial pressure, whether it comes from the Krogh-Erlang equation or the FEniCS solution, we draw a random number 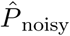 from a Gaussian distribution with mean 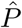 and standard deviation 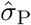, and replace 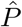 by this number.

### 2.2. Laplace estimator

Equation 5 says that given a data set of oxygen partial pressure 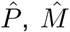 can be estimated by taking the Laplacian of 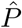. With dimensionless parameters we have

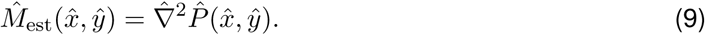

With 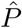 given on a square (or rectangular) grid with grid spacing 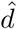, the net oxygen consumption as described by 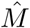 can be estimated at grid positions by using the discrete finite difference approximation of the Laplace operator:

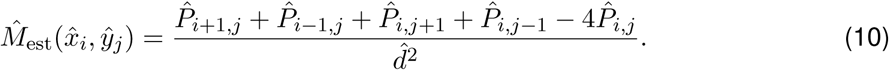

Here the integers *i* and *j* represent the grid positions, that is, 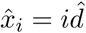 and 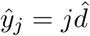.

In the present application, the MATLAB function `del2` is used to compute this discrete finite difference approximation of the Laplace operator. Note that in order to calculate the right-hand side of Equation 10, one must multiply the output from `del2` by 4. Specifically, we use the command 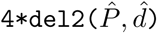 to calculate 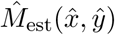.

#### 2.2.1. Smoothing

We reduce the adverse effects of noise in the oxygen pressure data by fitting a cubic smoothing spline to the 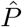 data before we calculate the Laplacian. Here smoothing is carried out using the `csaps` function in MATLAB’s Curve Fitting Toolbox. The `csaps` function takes a given data set 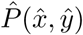 and generates a smoothing spline 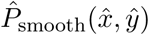 which minimizes

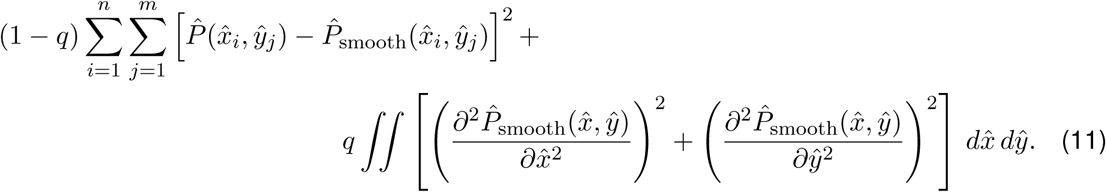

Here, *n* and *m* are the number of entries of 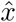 and *ŷ* respectively, and *q* is a smoothing parameter between 0 and 1. This smoothing routine penalizes large spatial double-derivatives in the estimated pressure 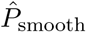 with the penalty parameterized by the parameter *q. q*=0 corresponds to the case with no smoothing, and increasing values of *q* imply increasing smoothing. Note that the `csaps` function in MATLAB takes *p* = 1 – *q* as input argument, see MATLAB documentation. This MATLAB function allows for giving more weights to some data points than others in the optimization. We keep the weights identical to 1 for all data points in the present application.

The `csaps` function allows the smoothing spline 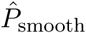 to be computed with higher resolution than the spatial resolution of the measurements. This is convenient as it allows for a higher spatial resolution in the maps of estimated *M* obtained from the discrete Laplace function `del2`. We here refer to the grid spacing between the pressure data points as 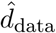, and the grid spacing of the estimated pressure points 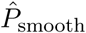 as 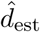. In the smoothing function, 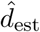 is set by inserting position vectors for the estimation points 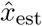 and *ŷ*_est_ with this spacing. Likewise, 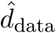 is set by inserting position vectors for the data points 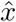 and *ŷ* with this spacing. Then 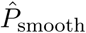 is estimated from the recorded pressure by the following call of `csaps`:

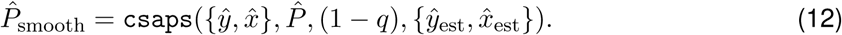

In the present paper we keep a fixed small value of 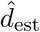, that is, 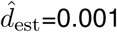. This value is set so small that the error introduced from the discreteness of the Laplace estimator in Equation 10 is negligible compared to other estimation errors.

#### 2.2.2. Choice of smoothing parameter

The effect of the `csaps` smoothing function can be characterized by a smoothing length 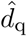 which describes how much a spatial *δ*-function is smeared out in space. By numerical exploration, we found that this characteristic smoothing length depends on *q* and 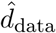 through the relationship

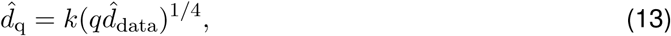

where *k* is a constant.

This relationship was found numerically by smoothing a square single-entry matrix with one as the center element, and the rest of the elements set to zero. The resulting spatially-smoothed *δ*-function was then plotted, for a fixed value of 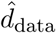 and different values of *q*, as a function of the distance *r* to the center point, as shown in Fig 1A. We then defined the characteristic length 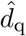 to be the distance from the center point at which the function value had fallen 50% compared to the center value, see dotted lines in panel A. Panel B shows the dependence of the estimated 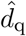 on *q* (for a fixed 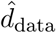 of 0.005). We observe that 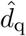 increases slowly with *q*, that is, when *q* is increased by a factor 10^4^, 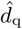 increases only by a factor 10. Fig 1C shows the smoothed *δ*-function when instead the value of *q* is fixed, while 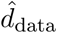 has different values. Again, when 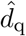 is read out from the curve and plotted as a function of 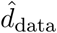 (panel D), we see that 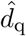 increases slowly with 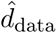, that is, when 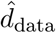 is increased by a factor 10^4^, 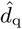 increases only by a factor 10.

**Figure 1:**
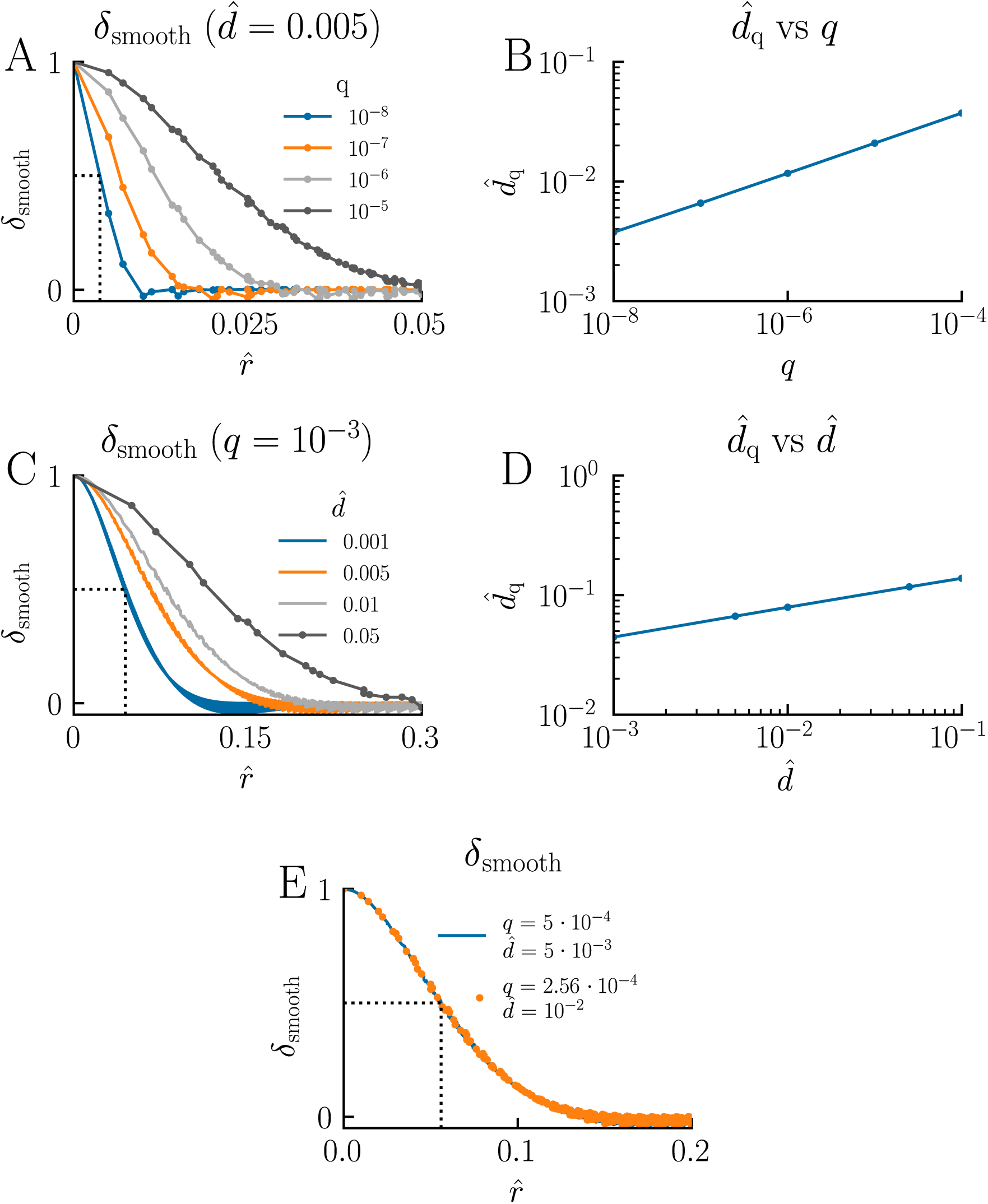
Choice of smoothing parameter in csaps. The effect of the smoothing function `csaps` is characterized by a smoothing length 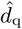 which is related to the smoothing factor *q* and the spatial spacing 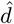 through Equation 13. We found this relationship by smoothing a two-dimensional spatial *δ*-function using different values of *q* and 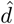, and plot the result as a function of the distance 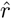 from the position of the *δ*-function. Panels A and C show the normalized smoothed *δ*-function 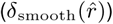 for different values of *q* (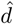 fixed) and 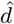 (*q* fixed), respectively. The characteristic smoothing length 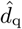 is defined as the distance corresponding to *δ*_smooth_ = 0.5 (dotted lines) and is plotted as a function of *q* and 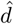 in panels B and D, respectively. In panel E we demonstrate how different sets of *q* and 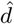-values correspond to the same 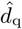, that is, the same smoothing effect.

The detailed value of the constant *k* in Equation 13 is not critical for our purpose. We set it by reading out the value for 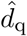 from the graph for the case with 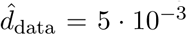 and *q* = 5 · 10^−4^ as shown with a blue line in Fig 1E. The readout value, 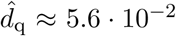, was then used to calculate *k* from Equation 13. After rounding to one decimal, this gave *k* = 1.4.

Thus, given 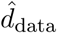 and a chosen value of 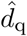, we can find which *q* to use in `csaps` in Equation 12 through the following formula:

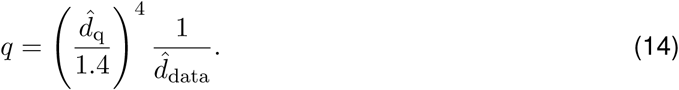

This equation tells us that if, say, 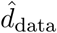 increases from 5 · 10^−3^ to 1 · 10^−2^, then *q* must decrease from *q* = 5 · 10^−4^ to about *q* = 2.6 · 10^−4^ to keep the same smoothing effect, that is, give the same value 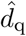. The dotted orange line in Fig 1E illustrates that this is indeed the case.

#### 2.2.3. Performance Measures of the Laplace Estimator

In order to evaluate the performance of the Laplace estimator, we test it on ground-truth data and calculate its bias, precision, and accuracy. As precision and accuracy measures we use standard deviation (SD) and root mean square error (RMSE). The mathematical definitions of these measures are

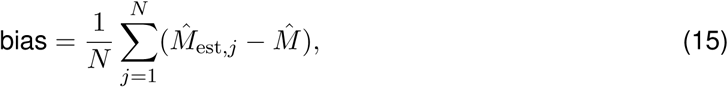

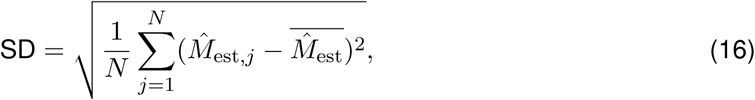

and

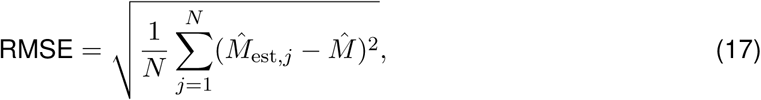

where *N* is the number of ground-truth samples and 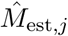 is the *j*^th^ estimate of 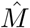.

The RMSE combines both bias and precision as its squared value MSE is equal to the standard deviation squared plus the bias squared: MSE = SD^2^ + bias^2^ [Wasserman, 2013].

## 3. Results

### 3.1. Illustration of Laplace estimation method

The principle of the Laplace method for estimation of the net oxygen consumption *M* (**r**) from measurements of the partial pressure *P* (**r**) of oxygen is illustrated in Fig 2. In this example we assume the spatial profile of the oxygen pressure to follow the Krogh-Erlang formula in Equation 6, mimicking the situation where a single arteriole is the source of the oxygen, and the oxygen consumption *M* is constant around the arteriole.

**Figure 2:**
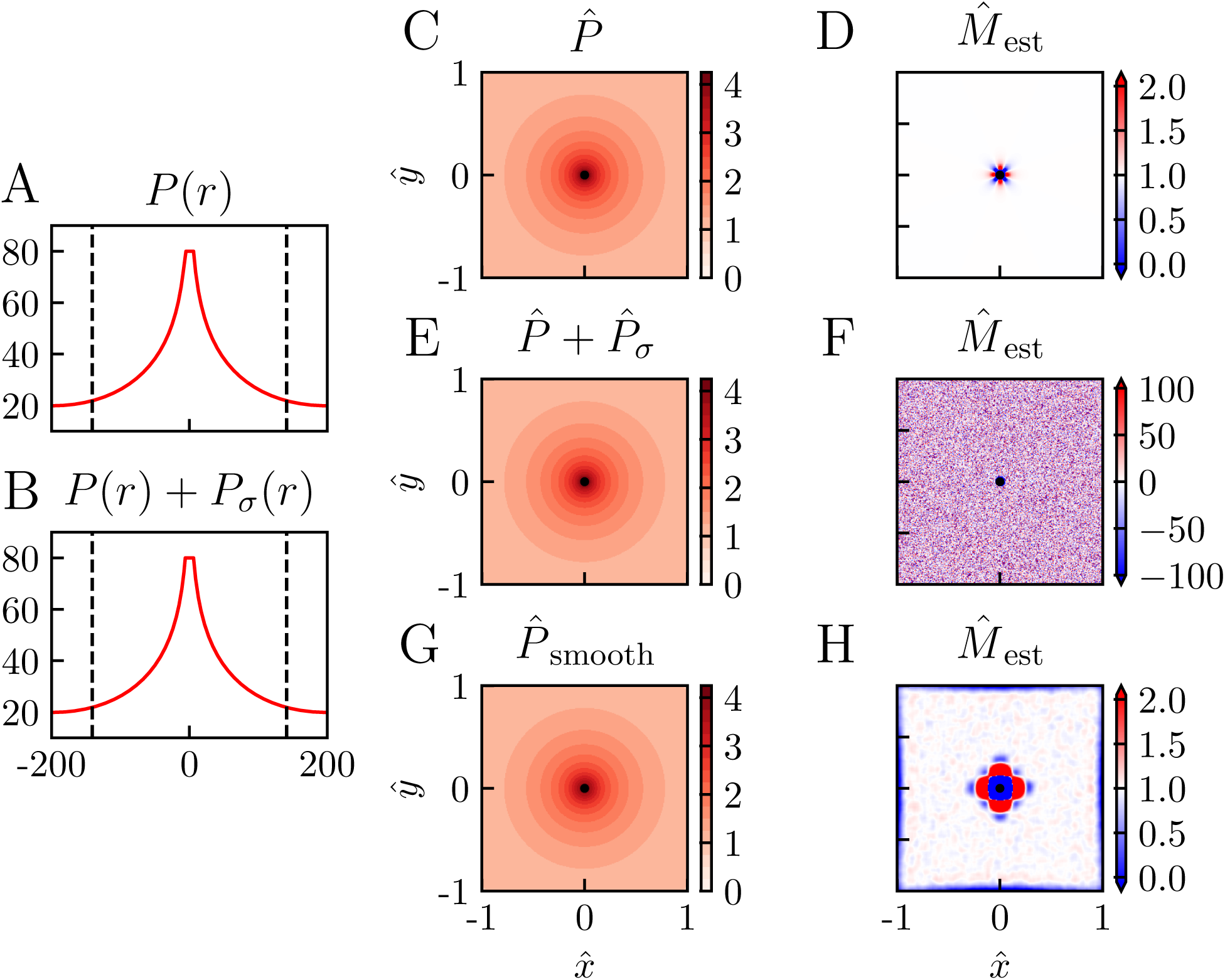
Illustration of Laplace estimation method. Panels A and B show examples of ground-truth pO2 profiles calculated using the Krogh-Erlang formula in Equation 7, with (panel A) and without noise (panel B). Panel C and E show the corresponding 2D representations of these pressure data sets, and panel G shows a data set where smoothing has been applied. Panel D, F and H show estimated *M* s calculated from the pO_2_ data in panel C, E and G, respectively. Parameter values: All panels: *P*_ves_ = 80 mmHg, *R*_ves_ = 6 *µ*m, *R*_t_ = 200 *µ*m, *M* = 10^−3^ mmHg*µ*m^−2^. For panels A,B: *d*_data_ = 1 *µ*m. For panel B: 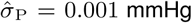. For panels C–H: 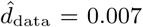, *r*^*^ = 141 *µ*m, *M* ^*^ = *M*. For panels E–H: 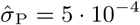. For panels G,H: 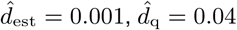.

Panel A shows the pressure profile in the radial directions as described by this formula with example parameters chosen to be in qualitative agreement with example data from Sakadžić et al. [2016], that is, *P*_ves_ = 80 mmHg, *M* = 10^−3^ mmHg*µ*m^−2^, *R*_ves_ = 6 *µ*m, and *R*_t_ = 200 *µ*m. Panel C shows a contour plot of this pressure profile in the two spatial dimensions. Here dimensionless parameters (cf. Methods) are used with *r*^*^ = 141 *µ*m and the convenient choice *M* ^*^ = *M* so that the maximal pressure *P*_ves_ corresponds to 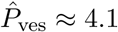 and 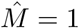. We show the pressure profile in a square window with side lenghts of 282 *µ*m so that the dimensionless position coordinates extends from −1 to 1 along the 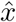 and *ŷ* axes. With this choice, the corners of the square correspond to a radial distance equal to 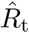, the radius of the tissue cylinder.

The problem of CMRO_2_ estimation now corresponds to estimating *M* at the different spatial positions inside the square window based on these recordings. Panel D shows the estimated *M* (in units of *M* ^*^) found by using the Laplace estimator in Equation 10 on the data in panel C. In this example the dimensionless distance between the grid points at which the pressure is ‘recorded’ is set to 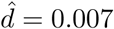, corresponding to a physical grid-point distance of about 1 *µm*. It is seen that some distance away from the vessel, the estimator predicts 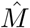 very close to 1, that is, *M* ≃ *M* ^*^, as it should.

However, close to the vessel, that is, for 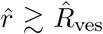, clearly incorrect values of 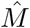 are estimated. One obvious reason is that the discrete Laplace estimator in Equation 10 will be inaccurate when one or more of the grid points used in the estimation is inside the vessel. Here the pressure *P* is not described by Equation 6 and is instead assumed constant so that ∇^2^*P* ≠ *M*, cf. Equation 3. For the present example a more important reason is that immediately outside the vessel, the pressure profile drops sharply (due to the last term in the Krogh-Erlang formula in Equation 6) so that the discrete Laplace estimator becomes inaccurate when the grid-point distance 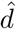 is too large. The ‘flower-like’ symmetric pattern of this estimation error in panel D reflects the cartesian symmetry of the estimator in Equation 10. This discretization error can be reduced by reducing the value of 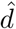.

Panel D in Fig 2 illustrates that if the experimental recordings were noiseless, the Laplace estimator in Equation 10 could be used directly on the oxygen pressure data, at least if the grid of recordings are finely spaced. This would apply for any distribution of vessels as long as the estimator 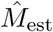 in Equation 10 is used sufficiently far away from oxygen-delivering blood vessels. Experimental pressure data will always contain noise, however, and panel B shows the pressure profile when an additive Gaussian noise *P*_*σ*_ with zero mean and standard deviation *σ*_P_ = 0.001 mmHg is added to the pressure signal in panel A. When 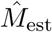 in Equation 10 is applied on the dimensionless version of these data (panel E), the estimated values of 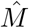 are wildly inaccurate (panel F). Not only does the estimated values of 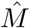 have much larger magnitudes than the true value of 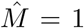, they also have both signs and vary strongly between neighboring grid positions (that is, between neighboring pixels in the panel image). These poor estimates reflect that the double-derivative operation of the Laplacian estimator corresponds to a high-pass spatial filtering which effectively amplifies the effects of the noise in the pressure recordings.

The high-frequency noise in the estimated 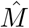 can be reduced by the use of spatial smoothing, that is, low-pass filtering, of the pressure data 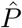 prior to application of 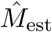. While the smoothed pressure profile 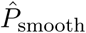 in panel G at first glance does not appear to be very different from the unsmoothed pressure in panel E, the effect of the smoothing on the estimated *M* is dramatic (panel H). With the choice of smoothing used in this example (see figure caption for details), quite accurate estimates of 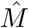 are found for a large region of the area around the central vessel (light-colored regions of panel H). However, the smoothing procedure results in large estimation errors in a sizable region around the blood vessel as well as close to the edges of the square data set.

As illustrated in this section, suitable smoothing of the oxygen partial pressure data before using the Laplace estimator 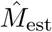 may dramatically improve the estimation accuracy. However, the choice of smoothing is critical: too little low-pass smoothing will not remove enough of the high-frequency spatial noise, too much smoothing will smooth away spatial information in the pressure signal and give poor estimates of *M* for this reason. Next, we will investigate this dilemma in more detail.

### 3.2. Noise removal vs. bias

Fig 3 illustrates the dilemma when choosing the right level of low-pass smoothing of the oxygen pressure data *P* before using the Laplace estimator in Equation 10. In the smoothing, the quantity described in Equation 11 was minimized to penalize sharp variations in *P*_smooth_ while at the same time fitting the ‘experimental’ data *P*. The level of smoothing is set by the smoothing length *d*_q_ (or 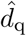 in dimensionless units) which is related to the smoothing parameter used in the presently used MATLAB function `csaps` via Equation 13 in Methods. This smoothing length describes how much a point (that is, a two-dimensional spatial *δ*-function) will be smeared out in space. Thus the larger *d*_q_ is, the more the pressure profile will be smeared out or smoothed.

**Figure 3:**
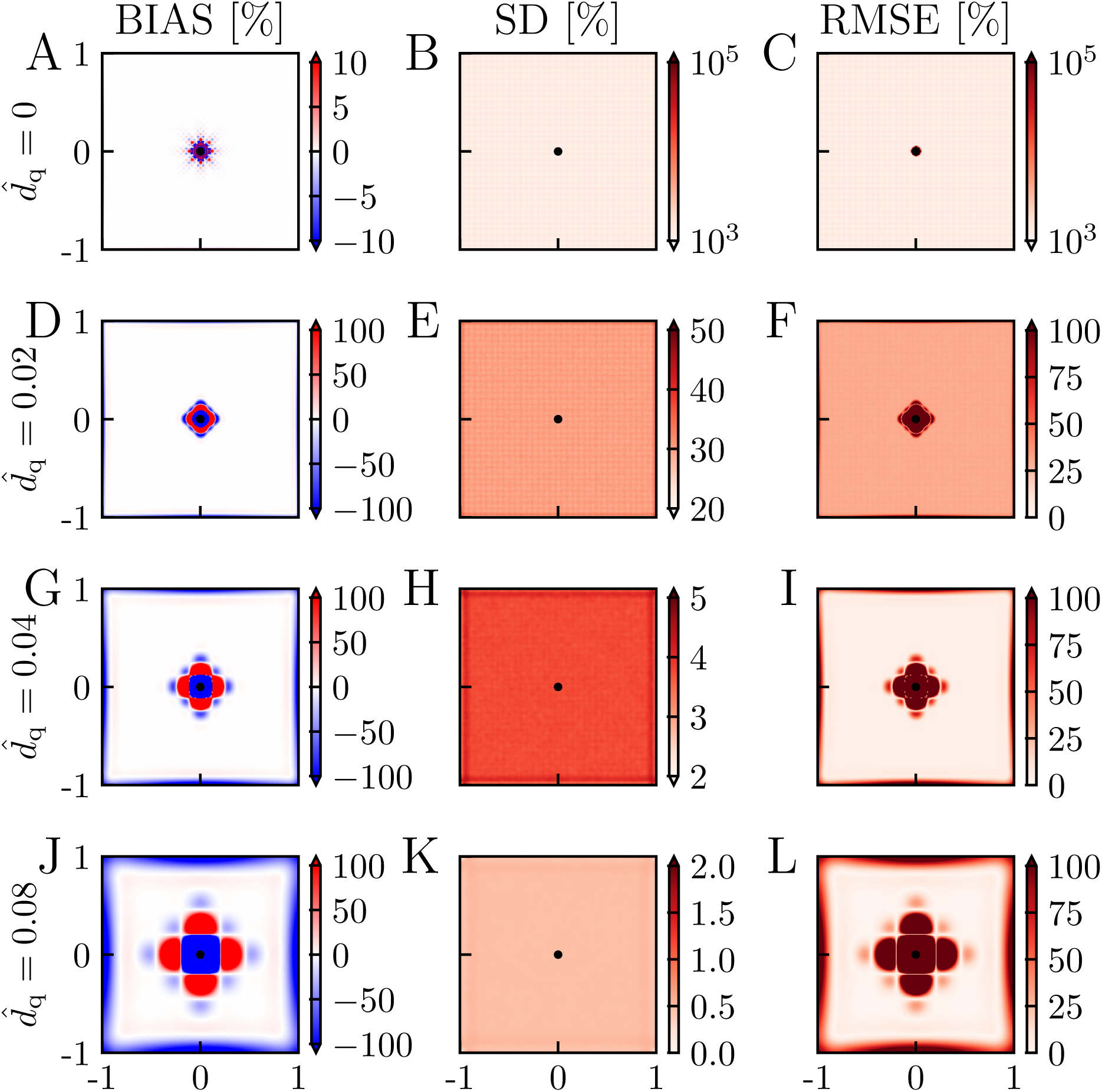
Illustration of noise removal vs bias. Bias is computed from Equation 15 for the case without noise 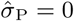 so that a single estimate of 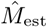 is sufficient, that is *N* =1 in Equation 15. SD is computed from Equation 16 with 10^4^ estimates of 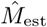, that is, *N* = 10^4^. In the computation of SD and RMSE,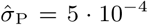. All performance measures are given as the percentage of the ground truth value 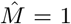. Note also that the MATLAB routine `csaps` is used also for the case without smoothing 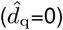 with *q*=0 inserted in Equation 12. Other parameter values: 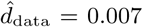, *P*_ves_ = 80 mmHg, *R*_ves_ = 6 *µ*m, *R*_t_ = 200 *µ*m, *M* = 10^−3^ mmHg*µ*m^−2^, *r*^*^ = 141 *µ*m, *M* ^*^ = *M*.

To quantify the performance of the estimator we use the three performance measures *bias, standard deviation (SD)*, and *root mean square error (RMSE)*. The bias (Equation 15) measures the systematic error in the estimator *M*_est_ introduced by the smoothing (and discreteness of data points) whether the data is noisy or not. It can be evaluated from noiseless data (that is, with *P*_*σ*_=0), and the results for different values of smoothing are shown in the panels in the left column of Fig 3 (panels A, D, G, J). In the case of no smoothing (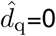, panel A) the only bias comes from the discreteness of the grid of data points, and a non-zero bias is only observed close to the vessel. With a small amount of smoothing (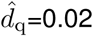, panel D), the bias around the vessel is increased. For 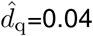 (panel G) and 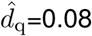 (panel J) this tendency of increased bias with increasing 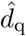 is continued, and some bias is also observed close to the edges of the square. For the largest smoothing depicted in panel J, about one-third or so of the estimation square has a bias with a magnitude larger than 100%.

The standard deviation (SD, Equation 16) measures the precision or the error in the estimation due to the presence of noise. This measure obviously depends on the level of noise *P*_*σ*_, and in the present example in Fig 3 a Gaussian noise with a standard deviation of 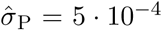 is used. With *r*^*^=141 *µ*m and *M* ^*^ = 10^−3^ mmHg*/µ*m^2^ as in Fig 2 this corresponds to a noise level of *σ*_P_ ≈ 0.01 mmHg. The SD for different amounts of smoothing is shown in the middle column of Fig 3 (panels B, E, H, K). Three observations of note are that (i) the SD of the estimates is extremely large when no smoothing is applied 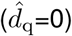, (ii) the SD decreases with increasing 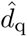, and (iii) unlike for the bias, the SD has similar values at the different positions.

An essential feature of the SD is that it is proportional to the standard deviation of the noise in the pressure 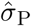. Thus if 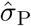 was doubled to 0.001, the SDs in panels B, E, and H would be doubled as well.

The accuracy of the estimator *M*_est_ is measured by the root mean square error (RMSE, Equation 17) which incorporates both the bias and the precision (SD) through the relation

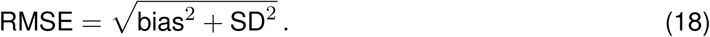

This measure describes the total statistical uncertainty of the estimates when *M*_est_ is applied on individual data sets. The bias increases with increasing 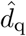 (panels A, D, G, J) while the SD instead decreases with increasing 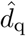 (panels B, E, H, K). One would thus expect a suitable intermediate value of 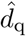 to give the smallest RMSE. For the example in Fig 3 we indeed see that of the values of 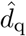 considered, the intermediate value 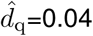 (panel I) offers the best compromise between bias and noise removal and gives the smallest RMSE. For this value of 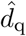 the RMSE is smaller than 25% for almost all positions except for a region around the blood vessel.

The large RMSE close to the blood vessel even for the ‘best’ choice of 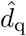 in panel I reflects the large bias in these positions (panel G).

### 3.3. Choice of smoothing length d_q_

As illustrated in the previous section, a key question when using the Laplace estimator in Equation 10 is the choice of the level of smoothing, or more specifically, the choice of the smoothing length *d*_q_. This will not only depend on the noise level, but also the spatial resolution of the experimental pressure data as described by the grid-point distance *d*_data_. Since the bias is independent of the noise level, and the SD is linearly proportional to the standard deviation *σ*_P_ of the noise, it is convenient to first explore the interplay between *d*_q_ and *d*_data_ for the bias and SD separately.

In Fig 4 we show how the bias varies with *d*_data_ and *d*_q_ for three choices of parameter values of each: 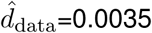, 0.007, 0.014 (here corresponding to physical grid-point distances of approximately 0.5 *µ*m, 1 *µ*m, and 2 *µ*m, respectively), 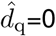, 0.02, 0.04 (corresponding to physical smoothing lengths of approximately 0 *µ*m, 3 *µ*m, and 6 *µ*m, respectively). For the case with no smoothing (panels A, D, G), we observe that the bias increases with increasing 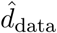. This illustrates that the error due to the discreteness of the Laplace estimator is sensitive to *d*_data_ even when *d*_est_ is set to a very small number (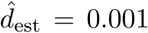, cf. Methods). This is not surprising because decreasing the grid-point distances from 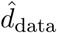 to 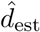 means that we estimate 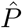 at a denser grid of points than what is directly available in the data. With smoothing added (two rightmost columns of panels), the bias increases, and the larger the value of 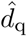, the larger the bias. (Note the difference in color scales in figure.)

**Figure 4:**
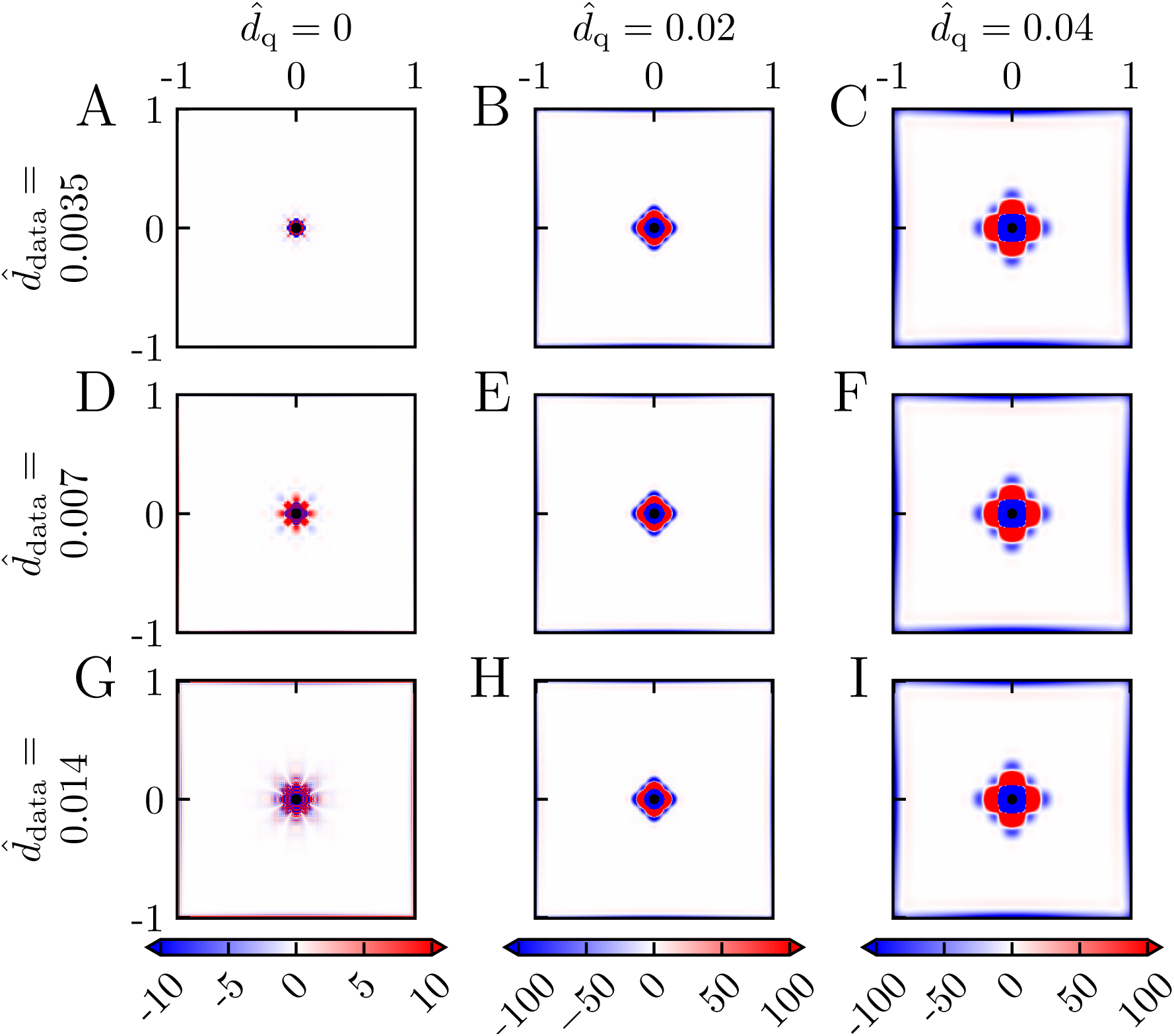
Bias for different smoothing. Bias computed from Equation 15 and given as the percentage of the ground truth value 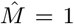. There was no noise added to the pressure data so that a single estimate of 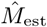 is sufficient, that is *N* =1 in Equation 15. Parameter values: *P*_ves_ = 80 mmHg, *R*_ves_ = 6 *µ*m, *R*_t_ = 200 *µ*m, *M* = 10^−3^ mmHg*µ*m^−2^, *r*^*^ = 141 *µ*m, *M* ^*^ = *M*.

In Fig 5 we correspondingly show how the SD (standard deviation) varies with *d*_data_ and *d*_q_ for the same set of parameters as in Fig 4 for a fixed level of noise in the pressure data 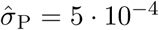. Here the most important feature is that the SD is strongly reduced with increased smoothing, that is, increasing *d*_q_ (from left to right). For the smoothed cases (two rightmost columns) we also observe that SD increases with increasing *d*_data_.

**Figure 5:**
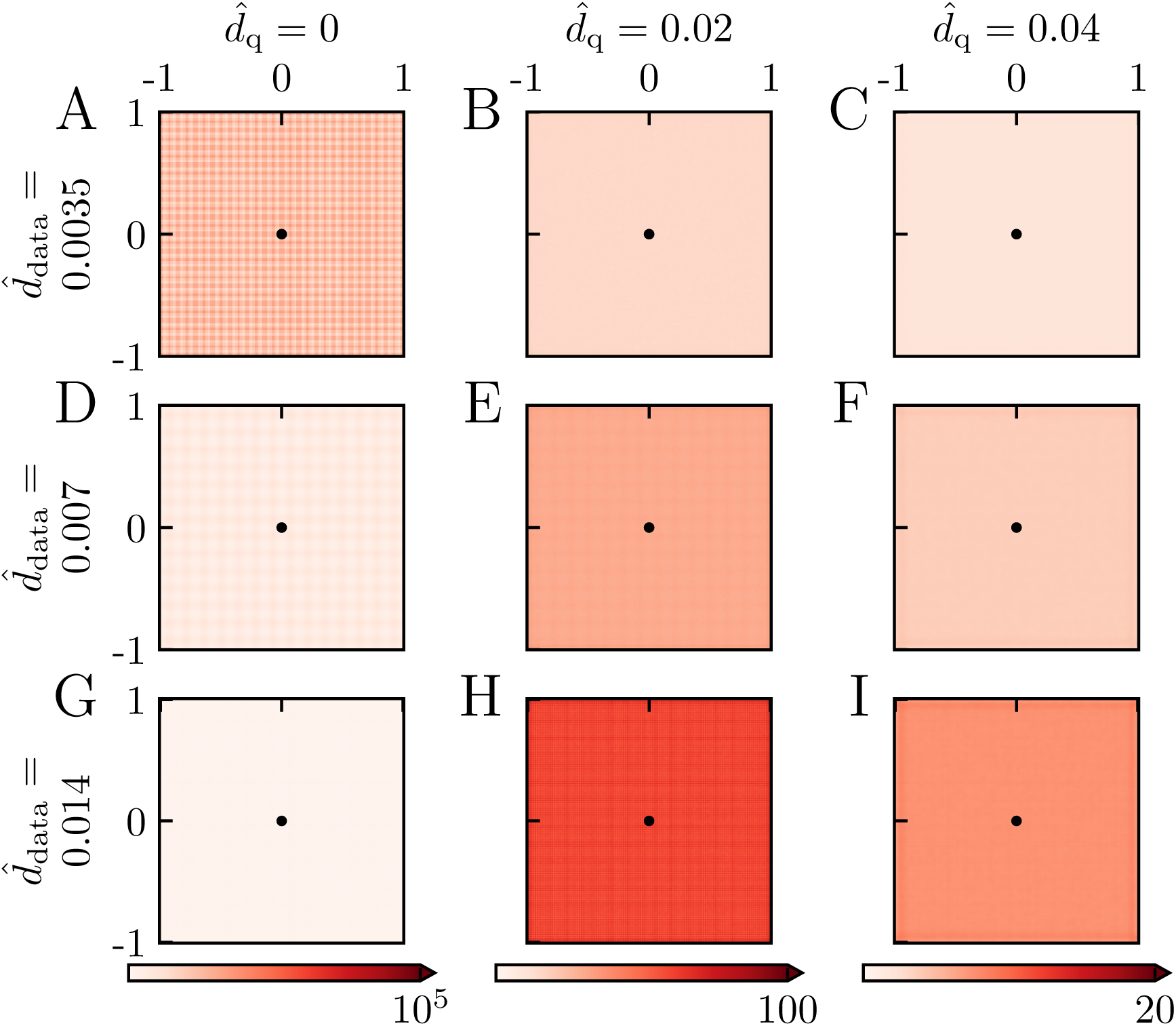
Standard deviation (SD) for different smoothing -fixed noise level. Standard deviation (SD) computed from Equation 16 with *N* = 10^4^. Values are given as the percentage of the ground truth value 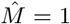. (Note that the grid-like pattern visible in some of the panels is a numerical artifact resulting from the application of the MATLAB routine `csaps`.) Parameter values: 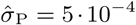, *P*_ves_ = 80 mmHg, *R*_ves_ = 6 *µ*m, *R*_t_ = 200 *µ*m, *M* = 10^−3^ mmHg*µ*m^−2^, *r*^*^ = 141 *µ*m, *M* ^*^ = *M*.

Fig 6 shows the RMSE, computed from Equation 18, for the example bias and SD shown in Fig 4 and Fig 5, respectively. For the smoothed cases (two right columns) we observe that the RMSE always increases with the 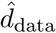. Thus with the noise level fixed, it is (unsurprisingly) always advantageous to have a dense recording grid. For the noise level in this example we see that the choice 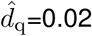 (second column) gives a good estimate for 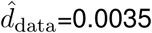, that is, low RMSE, for large parts of the estimation window. For 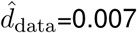 and especially 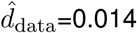 the SD is not sufficiently reduced, and the RMSE is overall high. For the case with a larger smoothing (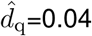, third column) the SD is much reduced for all values of 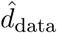. However, the region with large bias around the vessel is increased, and the spatial region in which small values of RMSE are shrunken.

**Figure 6:**
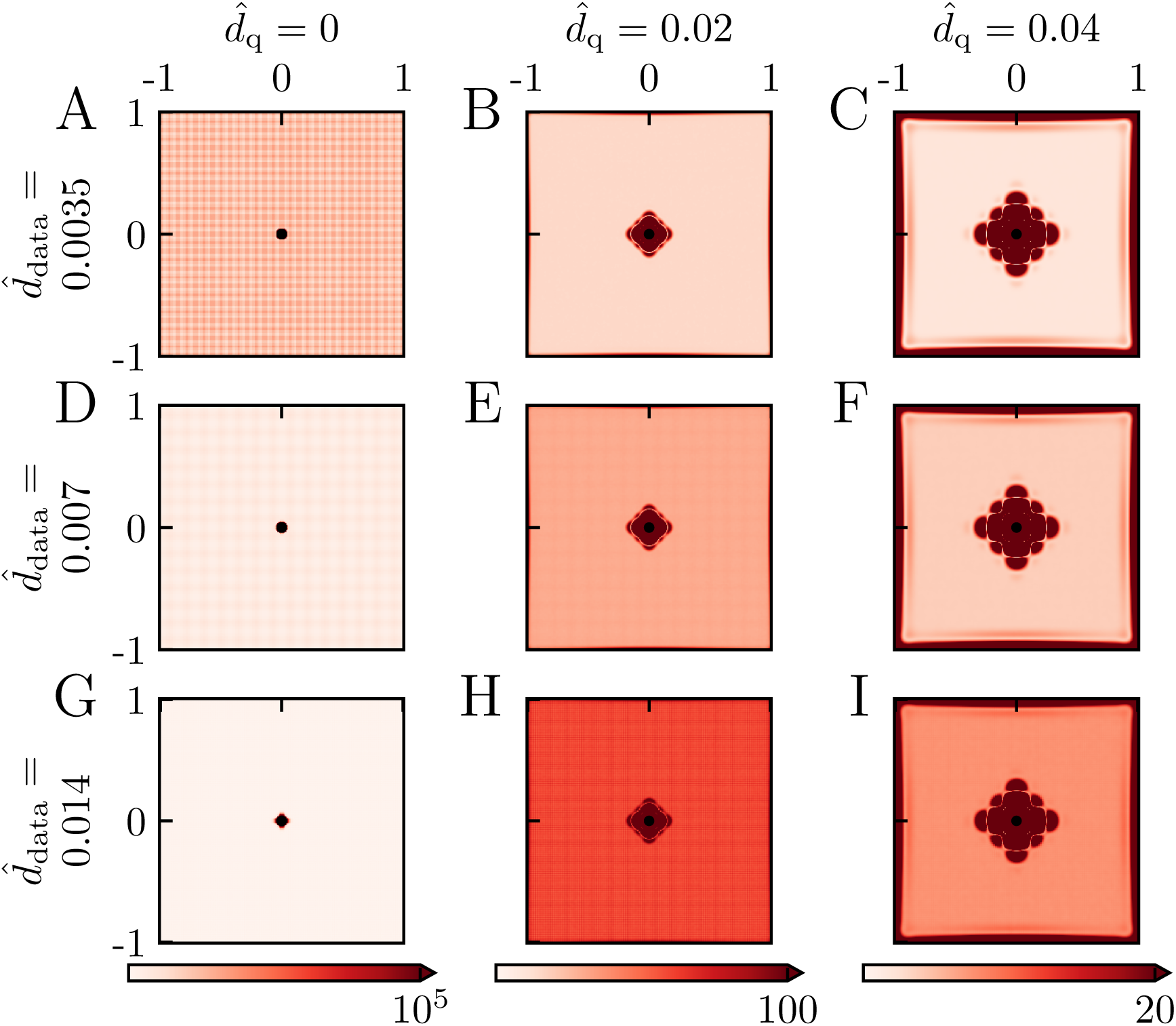
Root mean square error (RMSE) for different smoothing -fixed noise level. Root mean square error (RMSE) computed from Equation 17 for the bias and standard deviations (SD) shown in Figures 4 and 5, respectively. Values are given as the percentage of the ground truth value 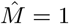.

Note that the SD results in Fig 5 and the RMSE results in Fig 6 only pertain to the particular noise level used in the example, that is, 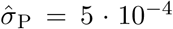. However, the SD is proportional to the noise level, so a doubling of 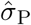 to 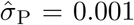 would simply double the SD from what is shown in Fig 5. RMSE results analogous to Fig 6 for other noise levels can thus be obtained by use of this SD scaling relationship in combination with Equation 18.

### 3.4. Estimation of CMRO_2_ *for other example situations*

In the examples above we have applied the Laplace estimator to the situation with (i) a constant value of the *M* and (ii) a single vessel providing the oxygen so that the pressure profile is described by the Krogh-Erlang formula in Equation 6. For these examples an alternative approach could be to estimate *M* using a ‘Krogh-estimator’, that is fitting the Krogh-Erlang formula directly to recorded data [Sakadžić et al., 2016]. In other situations where, for example, *M* varies with position or several nearby vessels provide the oxygen so that the circular symmetry assumed in the Krogh-Erlang formula is absent, such a Krogh estimator will not be applicable. However, the Laplace estimator does not assume a constant *M* or any particular arrangement of the oxygen-providing vessels and can be applied also here.

#### 3.4.1. Spatially varying CMRO_2_

To illustrate the applicability of the Laplace estimator to the situation with varying *M*, we consider in Fig 7 the situation where a single vessel provides the oxygen, but where the CMRO_2_ parameter *M* is four times larger in the upper half-plane than in the lower half-plane. Here the solution of the Poisson equation in Equation 3 must be found numerically, and in panel A we illustrate the oxygen pressure profile found using the FEniCS numerical solver (see Methods). While panel A shows the pressure profile without any added noise, panel B correspondingly shows a 2D colormap of the same pressure profile when noise has been added. In both panels we, as expected, observe that pressure is higher in the lower half-plane where the oxygen consumption, that is, *M*, is smallest so that the pressure profile decays more slowly with distance from the vessel.

**Figure 7:**
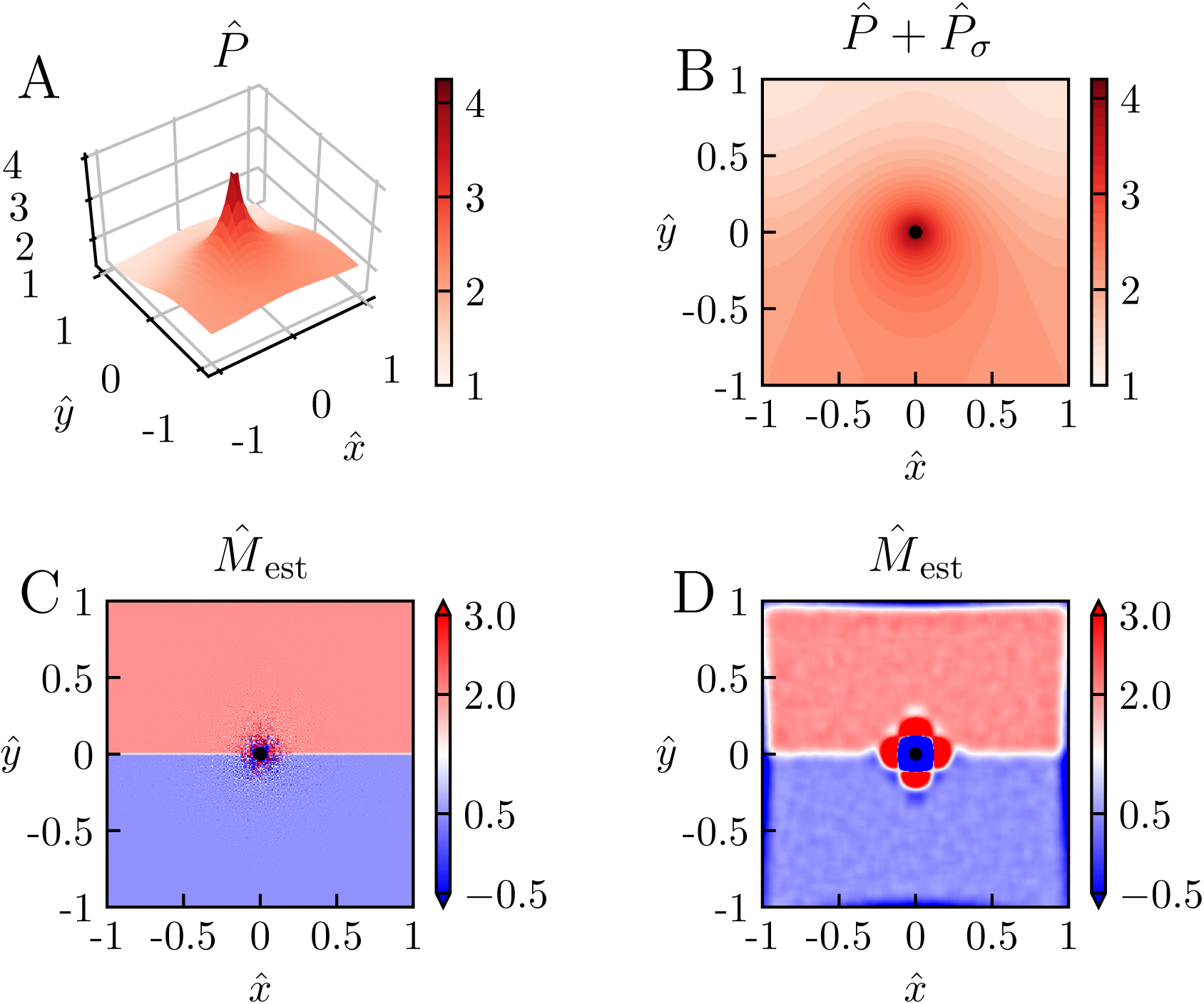
Estimation of *M* with spatially varying CMRO*2*. Laplace estimation of *M* for the case with a single oxygen-releasing vessel in the center with a larger CMRO_2_ in the upper half plane 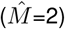 than in the lower half-plane 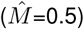. The ground-truth pO_2_ profile was calculated using the FEniCS numerical solver (see Methods). A: Illustration of pressure profile for the case without noise 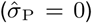. B: Illustration of pressure profile in A with noise added 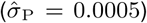. C: Estimated *M* from the noise-less profile in A without use of smoothing. D: Estimated *M* from the profile in B (where noise is present) with use of smoothing 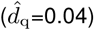. Other parameter values: 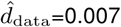, *P*_ves_ = 80 mmHg, *R*_ves_ = 6 *µ*m, *r*^*^=141 *µ*m, *M* ^*^ = 10^−3^.

When using the Laplace estimator on the noise-free data in Fig 7A, we obtain excellent estimates of *M*, that is, 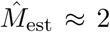 in the upper half-plane and 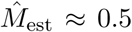 in the lower half-plane (panel C). We only observe sizable errors in the immediate vicinity of the vessel, errors stemming from the discreteness of the pressure data used in the estimation 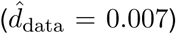. Further, when using the Laplace estimator on a smoothed version of the data in Fig 7B, we still obtain good estimates of 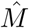 some distance away from the vessel. This is in accordance with the low values for the RMSE found for suitable smoothing of noisy pressure data for the case with constant 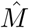 in Fig 6.

#### 3.4.2. Several vessels providing oxygen

An example of a situation where many nearby vessels contribute with oxygen is shown in Fig 8. Again no analytical solution of the pressure profile is available, and the Poisson equation is instead computed by means of FEniCS. As observed in the left panel, the circular symmetry of the pressure profile is broken around the vessel, but the Laplace estimator is still able to accurately estimate 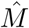 except close to the vessels (right panel).

**Figure 8:**
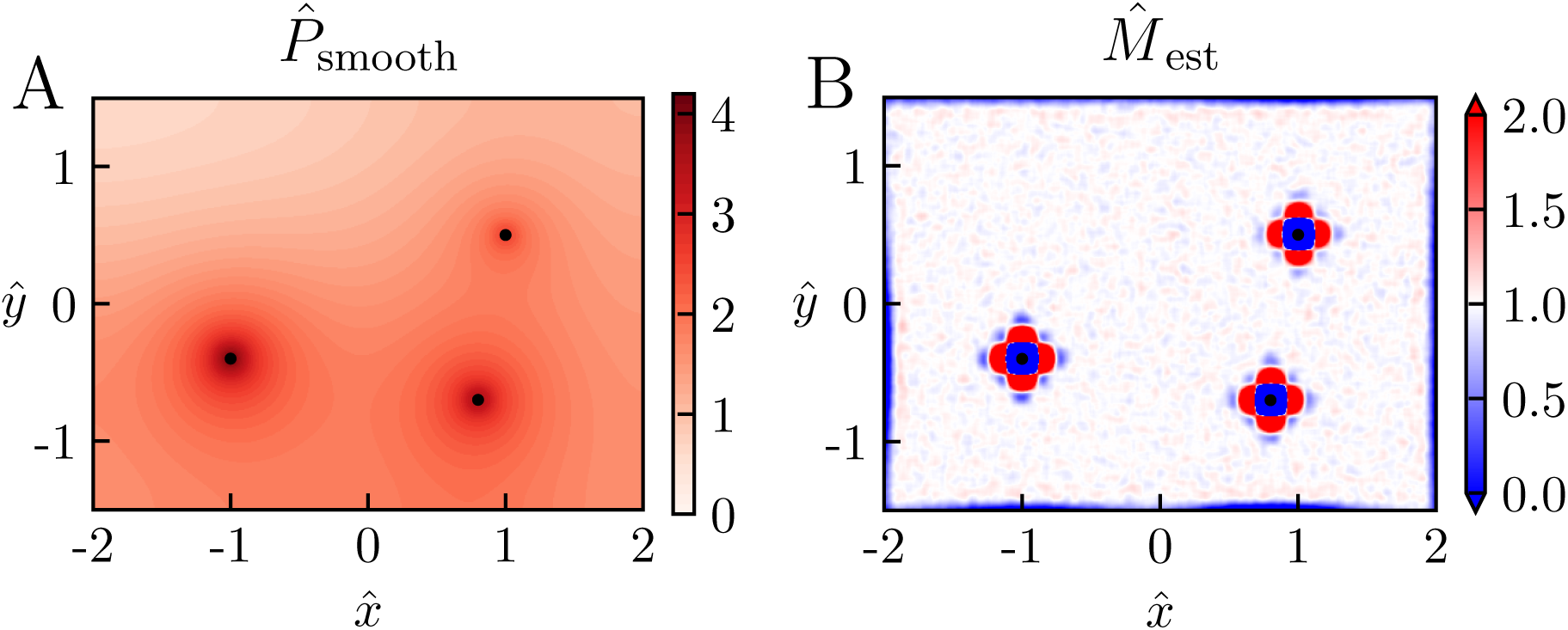
Estimation of *M* with several vessels providing oxygen. Example of Laplace estimation for a situation where three vessels release oxygen into the tissue. The ground-truth pO_2_ profile was calculated using the FEniCS numerical solver (see Methods). Here *P*_ves_ is set to 80 mmHg, 70 mmHg, and 50 mmHg for the vessel on the left, lower right, and upper right, respectively, while *R*_ves_ is set to 6 *µ*m for all vessels. Noise is added to the pressure profile in panel A 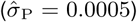, and 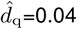 is used in the smoothing to provide the estimates of *M* in panel B. Other parameter values: 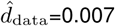, *M* = 10^−3^mmHg*µ*m^−2^, *r*^*^=141 *µ*m, *M* ^*^ = *M*.

### 3.5. Estimation of spatially-averaged M

So far we have used the Laplace estimator to estimate spatial maps of CMRO_2_ consumptions, that is, spatial maps of estimated *M*. The Laplace estimator can give accurate estimates as long as the noise level is not too large, but the estimates of *M* in the immediate vicinity of the oxygen-releasing blood vessels are typically inaccurate due to the bias introduced by the smoothing procedure.

In situations where the oxygen pressure data is too noisy to give reliable spatially resolved maps of estimated *M*, one can still obtain estimates of spatially-averaged values of *M* (as when estimating CMRO_2_ based on fitting the Krogh-Erlang model in Equation 6 to experimental data [Sakadžić et al., 2016]). The obvious procedure for estimating such average values *M*_est,av_ is to take the spatial average over spatially resolved values of *M*_est_, that is

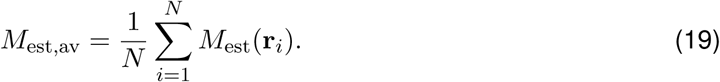

The SD of *M*_est,av_ is then expected to be a factor 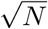 reduced compared to the SD for the spatially resolved estimates *M*_est_(**r**).

The bias is not reduced by such an averaging procedure, however. To reduce the effects of smoothing-induced bias, one possible procedure is to take the average of *M* only for positions outside a circular region around the oxygen-delivering vessel. As illustrated in Fig 9A this can reduce the bias in the *M*_est,av_ substantially. Larger values of the smoothing length 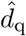 give larger regions of large bias around the vessel (Fig 4). Thus larger areas around the vessel, parameterized by the diameter 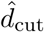, should be removed from the averaging sum in Equation 19 to keep the bias small. This removal of area from the averaging sum implies a smaller value for *N* in Equation 19 and thus a larger value of SD of *M*_est,av_. Again, a compromise between the bias and the SD must be found to get the most accurate estimate.

**Figure 9:**
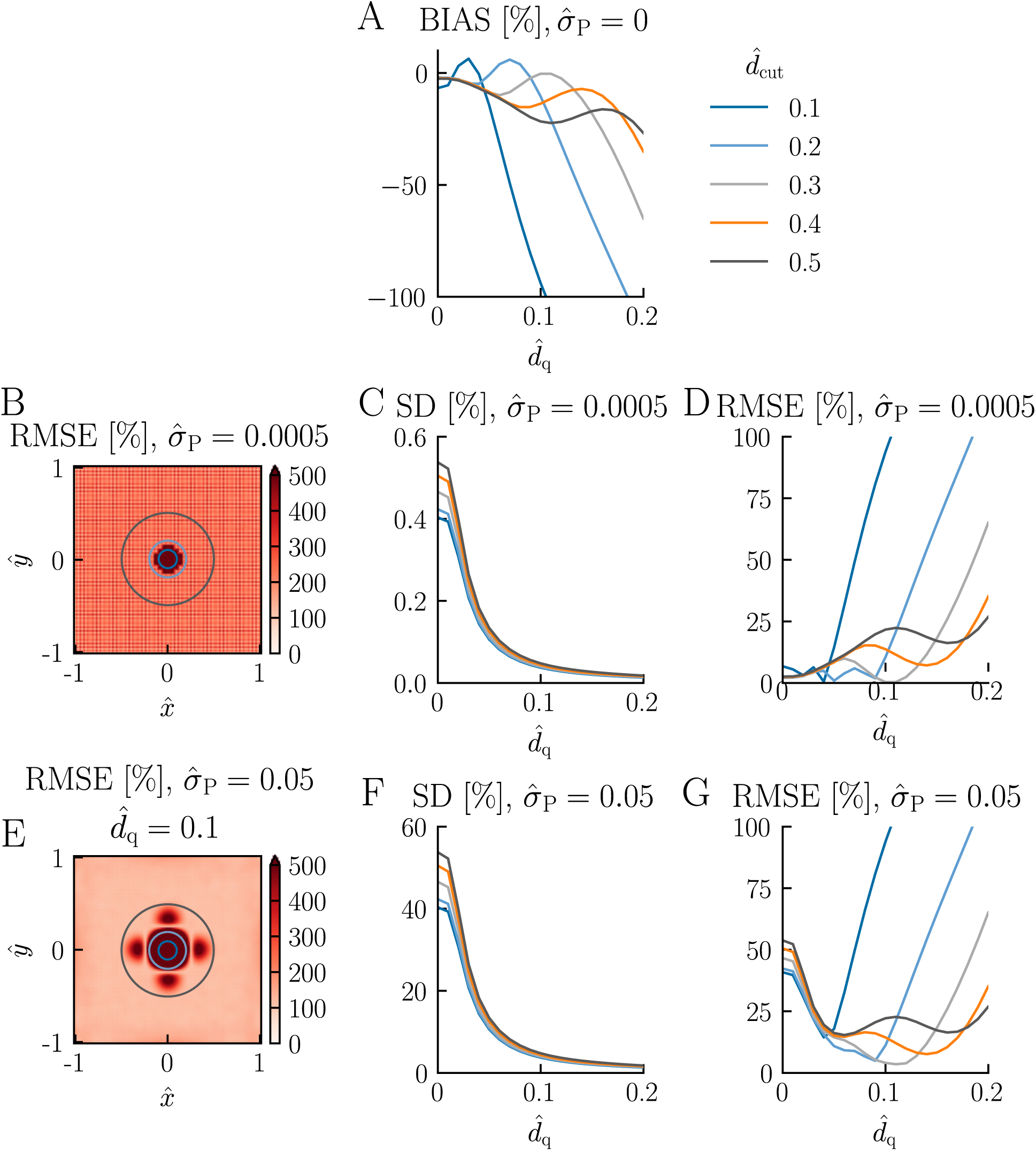
Estimation of spatially-averaged *M*. Illustration of accuracy of the estimation of spatially-averaged *M* for different values of the diameter 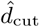 of the circular disc removed from the average in Equation 19. *N* = 1000 has been used in the estimation of the standard deviation (Equation 16). Other parameter values: All panels: 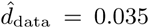, *P*_ves_ = 80 mmHg, *R*_ves_ = 6 *µ*m, *R*_t_ = 200 *µ*m, *M* = 10^−3^ mmHg*µ*m^−2^, *r*^*^ = 141 *µ*m, *M* ^*^ = *M*. For panel A: 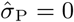. For panels B-D: 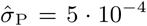. For panels E-G: 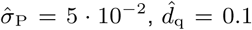. Note that for figure clarity, only the circles corresponding to 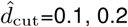, and 0.5 are shown in panels B and E.

This compromise is illustrated in Fig 9B–G. Panel B shows the spatially resolved RMSE for a case with low noise corresponding with no smoothing applied (cf. left column of Fig 6). Here the noise level is so low that even without smoothing, the SD of *M*_est,av_ becomes less than 1% for all averaging areas considered, that is, all choices of 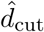 (cf. 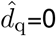 in panel C). With smoothing applied, the SD of *M*_est,av_ becomes even smaller, much less than 0.1% (panel C). We also note that the SD is largest for the largest value of 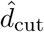, reflecting that here the averaging area (and thus *N* in Equation 19) is the smallest. The corresponding RMSE is shown in panel D. For this low-noise situation, there is nothing to gain by doing smoothing when estimating *M*_est,av_. The lowest RMSEs are obtained for 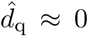 since smoothing reduces the accuracy of the estimates due to the bias introduced (cf. panel A).

The situation with a much higher noise level (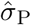 a factor 100 larger, that is,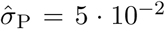) is shown in panels E–G. The spatially resolved RMSE using a smoothing factor of 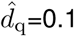 is seen to give large lobes with high RMSE values around the vessel (panel E). Moreover, the typical RMSE value outside the lobe region is about 120%. The SD of *M*_est,av_ (panel F) is seen to be on the order of 50% for the case without smoothing 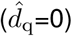, and a smaller RMSE can thus be obtained with smoothing applied (panel G). The smallest RMSE, less than ∼10%, is obtained for 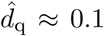 and 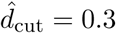.

This high-noise example illustrates how accurate estimates of *M*_av_ can be obtained even when the spatially resolved estimates for *M* have a large uncertainty. With the parameter values used here, that is, *M* ^*^ = 10^−3^ mmHg*µ*m^−2^ and *r*^*^ = 141 *µ*m, a 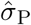 of 5 · 10^−2^ corresponds to a physical noise level *σ*_P_ of ≈ 1 mmHg. (Here we have used that 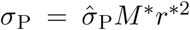, cf. Equation 5.) For comparison, the corresponding partial oxygen pressure at the vessel surface in this example would be *P*_ves_ = 80 mmHg.

## 4. Discussion

In the present paper we have introduced a new method, the *Laplace method*, to provide spatially resolved maps of CMRO_2_ estimates based on spatial measurements of pO_2_ [Sakadžić et al., 2010, 2016]. The method has two key steps: (i) spatial smoothing of measured pO_2_ profiles followed by (ii) application of double spatial derivatives in two spatial dimensions, that is, a Laplace operator. This method is an alternative to the Krogh-Erlang method where a spatially averaged value of CMRO_2_ is obtained around arterioles assuming circular symmetry [Sakadžić et al., 2016].

### 4.1. Improvement of Laplace method

The double spatial-derivative operation inherent in the Laplace approach is inherently sensitive to spatial noise, and the choice of a suitable smoothing method is thus essential for obtaining accurate CMRO_2_ estimates. The ideal smoothing method should reduce the effects of this spatial noise without introducing large biases in the resulting estimates.

Here we for convenience used the MATLAB smoothing function `csaps` which is publicly available and easy to use. `csaps` minimizes the functional in Equation 11 and thus penalizes large double spatial derivatives in the estimation of a smoothed pressure profile. Since CMRO_2_, or more precisely the variable *M* in Equation 3, is proportional to double spatial derivatives, this smoothing method effectively penalizes large magnitudes of *M* and thus introduces an unwanted bias. A better approach would have been to instead penalize changes in the spatial derivatives of *M*, that is, third spatial derivatives in the pressure. However, at present such a smoothing routine was not available to us.

While `csaps` allows for different weighting of different spatial positions in the smoothing process, the weighting functions are restricted to be spatially separable in the *x* and *y* directions. For the present application this limitation is not optimal as it would be preferable to include all positions except in a small region in and around the vessel, in the optimization inherent in the smoothing routine.

An obvious next step would thus be to test the accuracy of the Laplace method with a different smoothing method that (i) penalizes third spatial derivatives of the pressure and (ii) allows for arbitrary choices of weight functions in the functional to be minimized. In particular, it would be interesting to explore to what extent such a smoothing could reduce the size and magnitude of the lobes of large bias seen around the vessel center in Fig 4. The present MATLAB scripts, which can be found online at https://github.com/CINPLA/CMRO2estimation, are designed to allow for an easy change of smoothing method when they become available.

### 4.2. Use of Laplace method

Experimental data with less noise from better dyes and better acquisition systems will improve estimation accuracy [Sakadžić et al., 2016], but the accuracy of spatially resolved CMRO_2_ estimates will still be limited by the spatial noise of experimentally recorded pO_2_ profiles. Pooling of spatially-resolved estimates (as described in Equation 19) will always improve the accuracy, but this will be at the expense of spatial resolution. This trade-off can be investigated within the Laplace method using the scripts accompanying this paper. Estimation accuracy can be studied systematically with model-based ground-truth data (either based on the Krogh-Erlang model or based on FEniCS computations) using the same grid density and noise levels as in the experimental situation of interest.

## 5. Acknowledgements

Early versions of some of the present work have been published as a Master’s thesis [Sætra, 2016] and in abstract form [Sætra et al., 2017].

